# RealSeq2: a software integrated with UMI identification, error correction, and methylation modifications storing

**DOI:** 10.1101/2023.05.16.539668

**Authors:** Ke Wang, Mengmeng Song, Min Li, Tianyu Cui, Zhentian Liu, Enjie Yu, Huan Fang, Xuan Gao, Xuefeng Xia, Jiayin Wang, Yanfang Guan, Tao Liu, Xin Yi

## Abstract

High-throughput UMI technology sequencing is widely used in early tumor screening, detection, recurrence monitoring, etc. Detecting extremely low-frequency mutations is especially important for monitoring tumor recurrence, so high-precision data, as well as high-quality data, are required. We developed *RealSeq2*, a new integrated data-preprocessing software based on fastp and gencore, to achieve adapter removal, quality control, UMI identification, and generate consensus reads by clustering and error correction using multithreading in high-throughput next-generation sequencing background. *RealSeq2* also supports methylation data of 5-methylcytosine bisulfite-free sequencing. *RealSeq2* defined a new tag in SAM for storing methylation information, which is beneficial for co-identifying methylation sites and mutation sites for downstream analysis. *RealSeq2* includes three submodules: ReadsProfiler, ReadsCleaner, and ReadsRecycler. In addition, the output format file (BAM or SAM) is universal for downstream analyses.

*RealSeq2* is the preferred upstream analysis software for the co-detection of ultra-low frequency mutations and bisulfite-free methylation data. The error profile provides data support for downstream analysis. Additionally, XM tags will become a standard protocol for recording methylation signals.

## Introduction

High-accuracy downstream data analysis needs to obtain high-quality and high-confidence variations, which closely bound up quality control and the preprocessing of sequencing data. Several complicated influence factors may cause data distortion, for instance, adapter contamination, base content biases, and overrepresented sequence. Worse, errors in library preparation and sequencing procedures are inevitable, which lead to misjudgments about the original nucleic acid sequence. Aiming to optimize the identification of sequencing reads derived from different DNA fragments, a technology named unique molecular identifier (UMI) has been developed to reduce errors introduced by DNA polymerase and amplification during sequencing.

Except for the above factors affecting sequencing quality, data preprocessing of raw data was an extremely critical step for further analysis. Several relevant preprocessing tools: FastProNGS (Liu, et al., 2019), FASTQC (https://www.bioinformatics.babraham.ac.uk/projects/fastqc/), Cutadapt (Martin, 2011), and Trimmomatic (Bolger, et al., 2014) could be considered as the feature-cornerstones for fastp (Chen, et al., 2018), which performs quality control, reads filtering, and bases correction for FASTQ data with an ultra-fast speed (2–5 times faster than any of the three mentioned methods above). Moreover, fastp could also support UMI preprocessing, per-read polyG tail trimming, and output splitting. It is much faster than its peers. However, variable-length UMIs are incompatible with this software, which could cause the loss of a certain quantity of high-quality data input. Furthermore, one of the most essential elements for quality control and downstream analysis is a substantial statistical result, nevertheless, rarely implemented in existing tools. After data preprocessing, UMI-integrated data need to process by gencore (Chen, et al., 2019), which is an efficient and powerful tool for duplicate removal and sequence error suppressing applied in UMI technology (Chen, S. et al, 2017). However, it occupies too many computing resources while dealing with UMIs. Therefore, it is necessary to develop a high-speed software that combines data pre-processing and UMI data processing functions.

In recent years, a bisulfite-free methylation detection technology has been applied (Liu, et al., 2019), which makes up for the shortcomings of the traditional whole Genome Bisulfite Sequencing (WGBS) DNA high damage and low complexity library (Chatterjee, et al., 2012; Fraga and Esteller, 2002; Frommer, et al., 1992), which can directly detect modifications with high sensitivity and specificity without affecting unmodified cytosine. How to make full use of genome-wide data to achieve methylation and mutation co-detection is exactly the problem we need to solve.

In view of the above problems, we have developed *RealSeq2*, referring to fastp and gencore. Besides possessing the features of handling UMIs and reporting informative results, *RealSeq2* combined simplicity with speediness. For instance, it optimizes at the algorithmic level (innovative optimization using the Bayesian algorithm) to remove noise and support downstream analysis. Its particular data dealing methodology could support bisulfite-free data to guarantee the maximum methylation signal without losing the clustering error correction performance.

## Methods

### Improvement of RealSeq2

*RealSeq2* is a data-preprocessing software specially used for UMI data processing. It contains three sub-modules: ReadsProfiler, ReadsCleaner, and ReadsRecycler. The ReadsProfiler module is mainly based on fastp software for secondary development. On the basis of retaining the short running time of fastp, it also supports the identification and trimming of variable-length UMIs, allows 1bp errors (mismatch and deletion) in UMI sequences, and outputs the error profile of each sequencing cycle (Table 1). As the core module of UMI clustering and error correction, the ReadsCleaner module refers to the clustering error correction strategy of gencore and fgbio software (https://github.com/fulcrumgenomics/fgbio), which can accurately identify and correct the sequencing errors in the sequence using the Bayesian algorithm. At present, Tair (Liu, et al., 2019) and bismark (Krueger and Andrews, 2011) are specialized applied for processing DNA modification data. There is a lack of tools for simultaneous analysis of mutation and methylation. *RealSeq2* is such a tool (Table 2). Using the information of the forward and reverse templates, the bases changed due to methylation in the data are reverted to normal bases and recorded the methylation information of the whole reads, including methylation position and strand through introducing a new tag XM to simultaneously record the methylation signal of the forward and reverse strands. The third module of *RealSeq2, ReadsRecycler*, is specially designed for bisulfite-free methylation data. It is used to process sequences that have not been aligned to the reference genome due to a large number of base changes caused by methylation. ReadsRecycler could correct and realign these unmapped sequences to improve data usage. In summary, *RealSeq2* can meet the quality control requirements of FASTQ format data and clustering error correction requirements of BAM format data, and it is also compatible with the processing of bisulfite-free sequencing data. The entire processes from the raw data as input to the aligned BAM as output is shown in Figure 1.

**Table 1.** Function comparison of quality control software supporting UMI

| Software | UMI<br>trimming | Visual<br>report<br>for UMI<br>sequence | UMI<br>error<br>toleration | support variable-length<br>UMI |
| --- | --- | --- | --- | --- |
| *RealSeq2-*<br>*ReadsProfiler* | true | true | true | true |
| *fastp* | true | false | false | false |
| *cutadapt* | false | false | false | false |

**Table 2.**
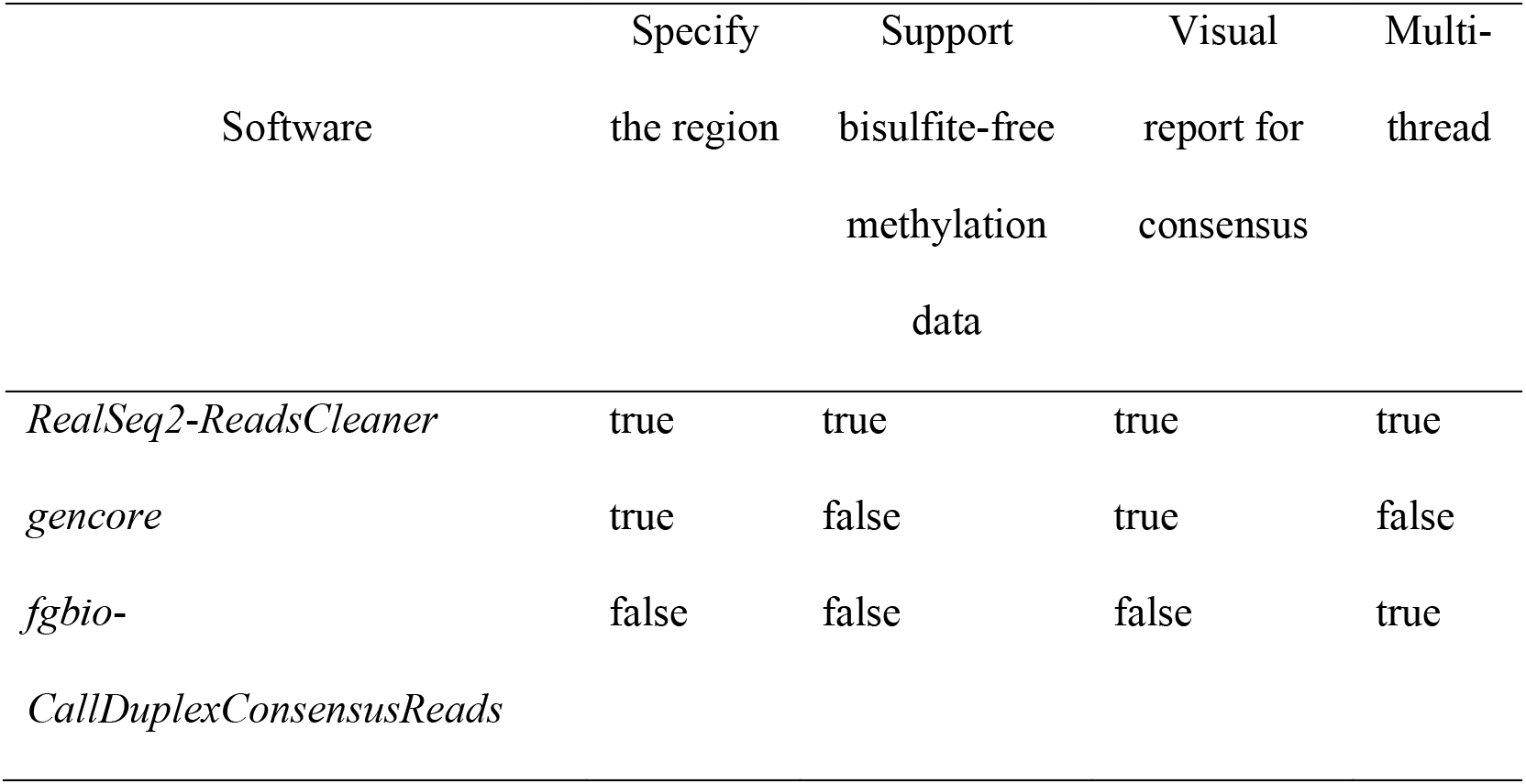
Function comparison of clustering error correction software

**Figure 1.**
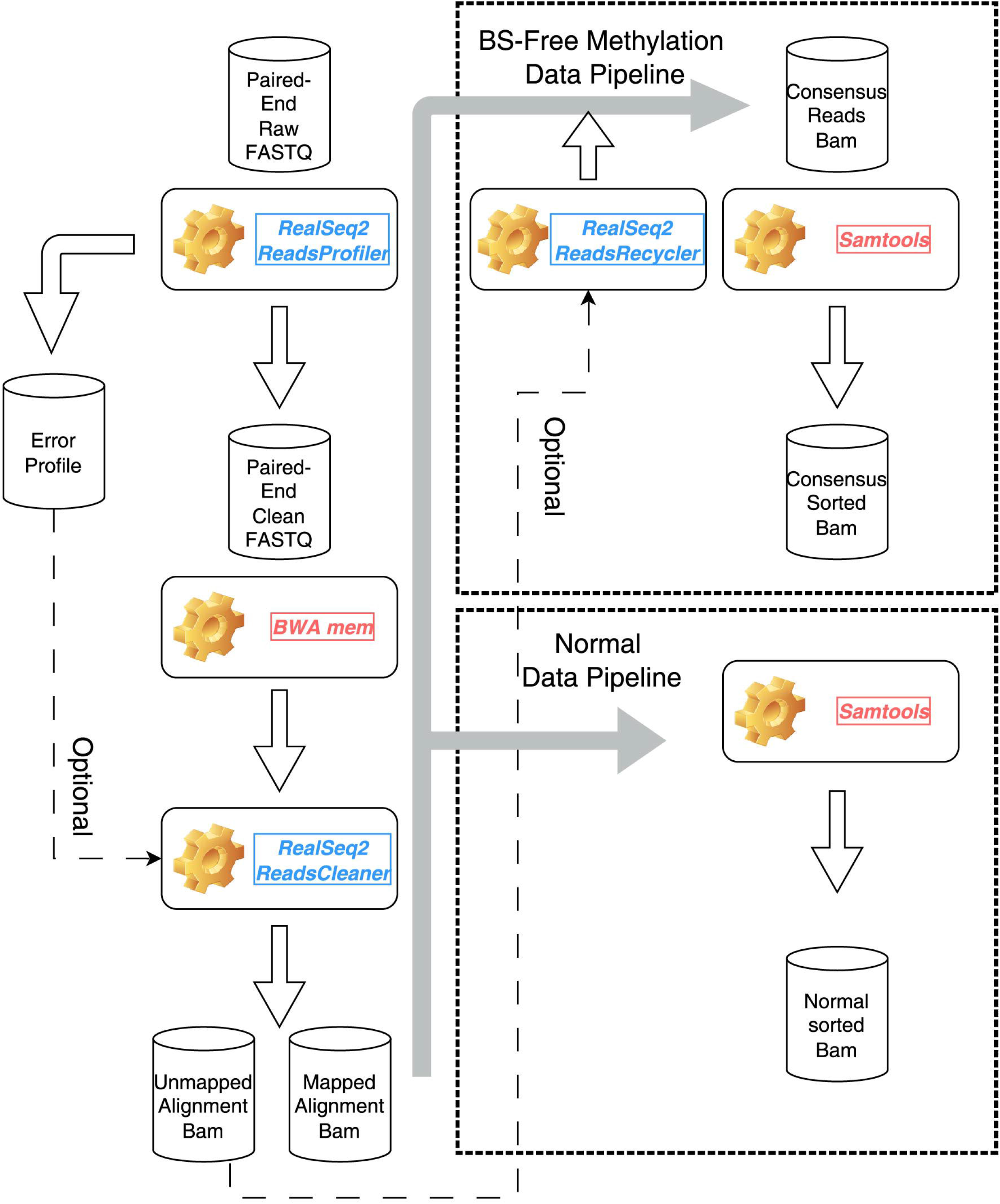
Full pipeline for RealSeq2.

### Identification of variable-length UMI

UMI identification is implemented by the ReadsProfiler submodule, which is reconstructed based on fastp software. It inherits quality control of raw FASTQ format files and adds functions of error correction of UMI sequence, variable-length UMI identification and trimming, and recording base error profile of each sequencing cycle. ReadsProfiler can identify variable-length UMIs and tolerant 1bp mismatch or deletion. Firstly, we binary-encode a set of known UMI sequences used in the library construction and all the fake UMI sequences for which 1bp mismatch or 1bp del may occur, and these results are built into a UMI binary code set. Secondly, the 5′ end of each read is matched to the UMI binary code set and preferentially matches the UMI sequence with the longest length in the UMI binary code set. Furthermore, if it matches exactly, the UMI sequence of the 5′ end read will be trimmed, and the UMI sequence will be recorded into the read name. If it does not exactly match, then it matches with the UMI with the 1bp error, trim the UMI sequence of the 5′ end reads, and record its corresponding real UMI with the read name. If the read matches none of the UMI sequences of that length, then matches with the next longest UMI sequences, loop through this step until finding a matching UMI sequence. The pseudocode for the UMI identification process is shown in Algorithm 1.

#### Algorithm 1 UMI identification and fault tolerance

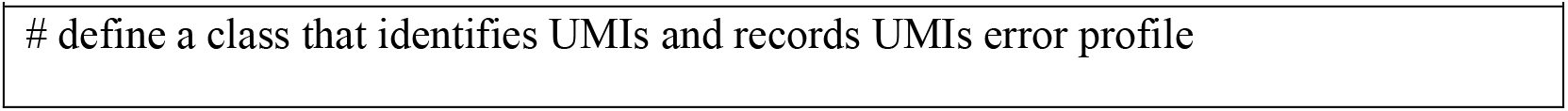
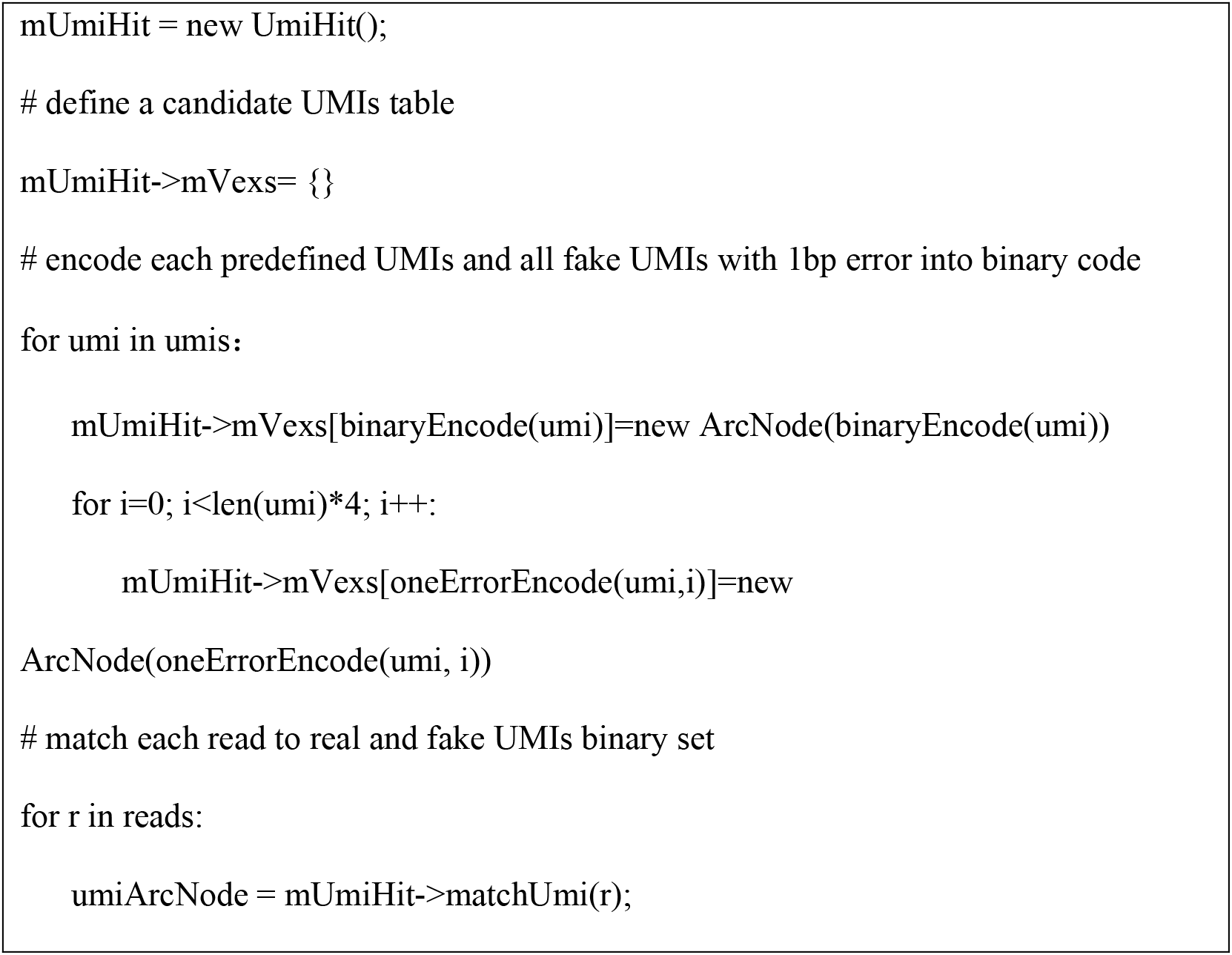

In addition, ReadsProfiler will also calculate the error rate of each sequencing cycle in the paired-end NGS UMI data at the same time as identifying UMIs. The principle is: UMIs are inserted into the start and end of each DNA fragment, respectively, and paired-end sequencing enables both ends of the DNA fragment to be sequenced with the forward read called read1 and the reverse read called read2. When the insert length is short, both read1 and read2 will detect each other’s UMIs based on the set of known UMI reference sequences. We identify and cluster the UMI sequences at the 5′ end of the sequencing data to generate consensus sequences and compare them with the 3′-end UMI of the corresponding reversed base sequence to identify possible sequencing errors at the 3′-end UMI. Counting the number of reads with this error at each location of the 3′-end UMI, calculating the error rate of each cycle 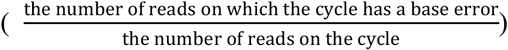, then the error profiles are saved in the Jason format file.

### Generate consensus reads based on UMI clusters

The ReadsCleaner submodule uses the “producer-consumer” multi-threaded concurrent cooperation pattern to cluster sequences based on the basial functions of gencore software. In this mode, all information read by a single thread will be temporarily stored in a fixed-size buffer, and simultaneously multiple consumer threads will read data from the buffer for processing. In the process of single-threaded reading sorted bam file, ReadsCleaner will first filter out unmapped sequences and mate unmapped sequences, and the mapped sequences will participate in the process of cluster error correction. ReadsCleaner puts the sequences with the same chromosome ID, alignment position, and mate alignment position into the same cluster, called “position cluster” and is the basic unit of subsequent multithreading processing. In the process of constructing a position cluster, ReadsCleaner first puts the sequence into the corresponding position cluster according to the currently read sequence alignment position. When the chromosome ID of a position cluster is inconsistent with the subsequent read sequence, or the alignment position is smaller than the position of the subsequent sequence alignment minus 1000bp, the position cluster is considered to have no new sequence. ReadsCleaner puts all position clusters that will not add new sequences into the buffer, waiting for the consumer thread to process.

When a position cluster enters the consumer thread, ReadsCleaner identifies the UMI tags of all sequences in the cluster. According to different UMI tags, all sequences are further classified into different UMI clusters. The basic unit of cluster error correction module processing is a UMI cluster. After error correction, each UMI cluster will only generate one consensus sequence (described in detail in **Methods section: Error correction of UMI cluster reads)**. Two consensus sequences from the same position cluster with complementary UMIs will be further duplex merged and finally achieve the purpose of removing PCR amplification errors and sequencing errors. Suppose the input is bisulfite-free library construction of 5-methylcytosine (bisulfite-free) methylation data (described in detail in section 2.4). In that case, *RealSeq2* will identify the methylation signal according to the bisulfite-free library principle during duplex merging (Figure 2). The principle is that one of the duplex sequences undergoes a base change from C to T or G to A (the bases at the current position of duplex sequences were the same in the templates which did not occur methylation modification at the current position), while the base at the same position of the other sequence is consistent with the reference genome, and *RealSeq2* considers that methylation modification occurs at this position. The base of the changed sequence will be corrected to be a base consistent with the reference genome and record the methylation signal in the SAM file.

**Figure 2.**
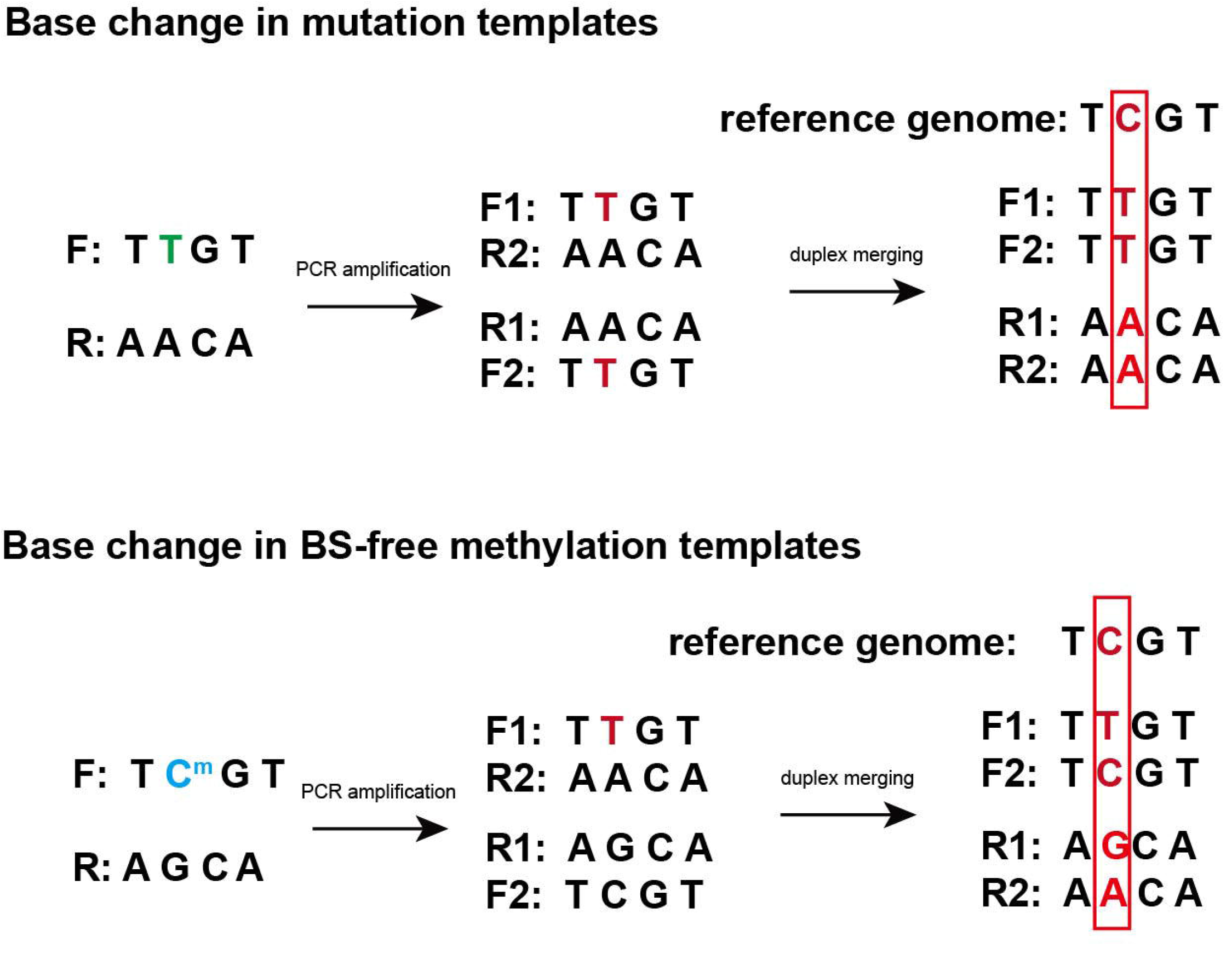
Diagram of base change of duplex sequences in bisulfite-free methylation sequencing.

### Error correction of UMI cluster reads

The schematic diagram of sequence clustering error correction is shown in Figure 3. The error correction process is the core part of the ReadsCleaner sub-module and adopts the principle of Bayesian posterior probability. After ReadsProfiler performs quality control on the data and UMI identification, it will output the error profile. Since the error profile is the result of the error rate of all sequences, and each base quality value is base-specific, in the ReadsCleaner algorithm, the error profile and base quality values are divided into two steps involved in the calculation.

**Figure 3.**
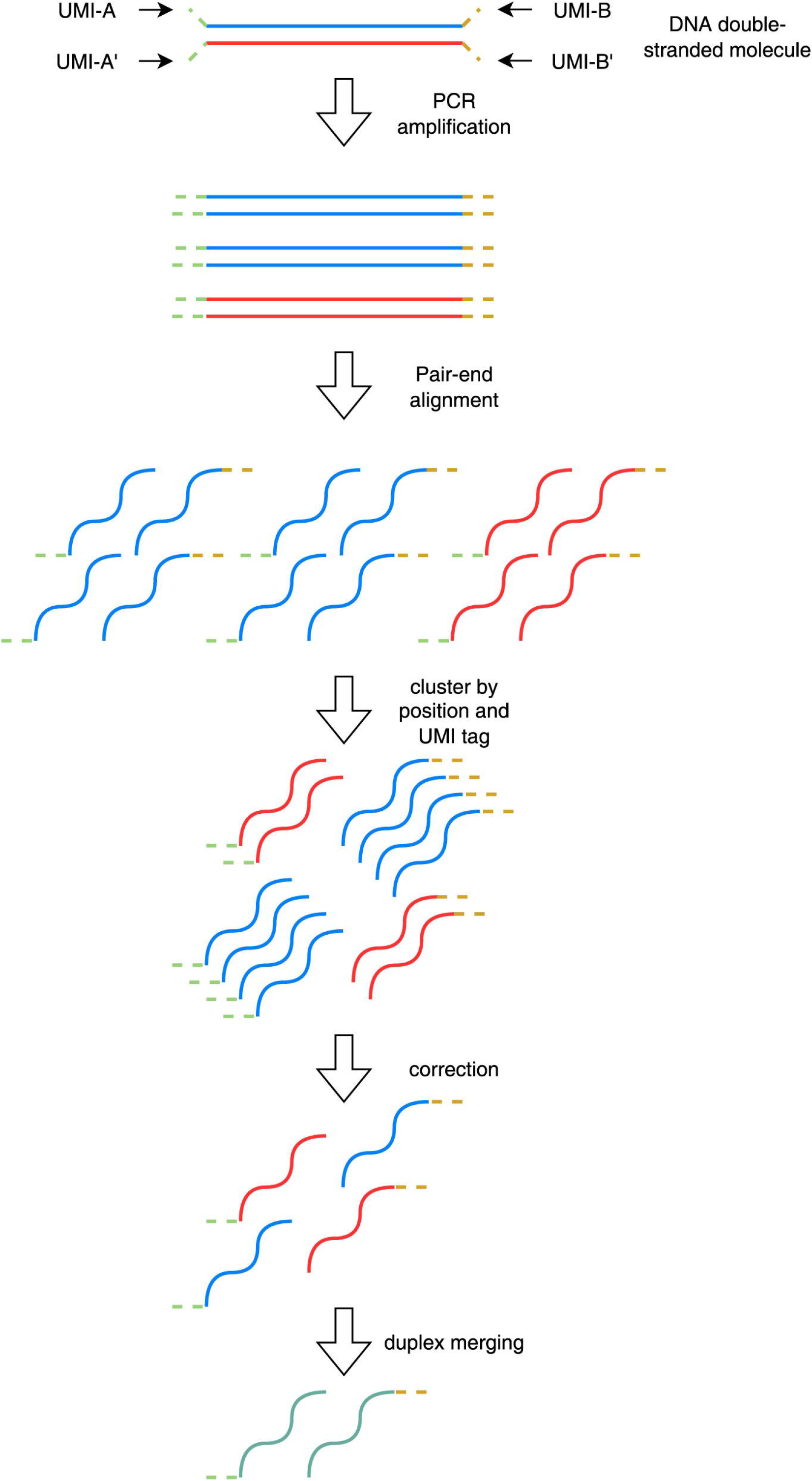
Diagram of reads clustering and error correction.

First, ReadsCleaner uses the error profile to correct the base quality value of the sequence, so that the base quality value not only reflects the base sequencing situation but also introduces the real sequencing error rate of the entire data. The final corrected base quality value represents the probability of an error. Since the quality value of each base is independent of each other, after correcting the base quality value, ReadsCleaner will cycle through all cycles, use the method of maximum posterior probability to calculate the base form of each cycle on the sequence, and the posterior probability of the base is used as the base quality value. The specific algorithm is as follows:

The sequencing length is represented by *k*, the number of total sequences in the UMI cluster is represented by *N*, the sequencing base of the *l*-th cycle of the *i*-th sequence is *base*_*i,l*_ with the base quality score *qual*_*i,j*_, the real base of the *l*-th cycle of the *i*-th sequence is *cns*_*i,l*_ We suppose the consensus sequence is known, the error probability *cycleErr*_*l,X* − >*Y*_ of *X* → *Y* in the *l*-th cycle can be expressed by formula (1):

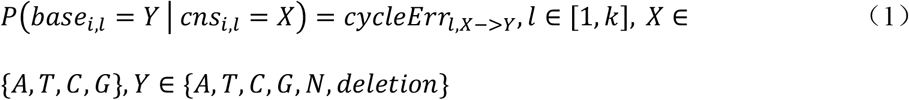

According to the Bayesian formula transformation, the error probability *cycleErr*_*l,X* − >*Y*_ can be converted into the probability of the current base occurred error in the sequencing, as shown in formula (2):

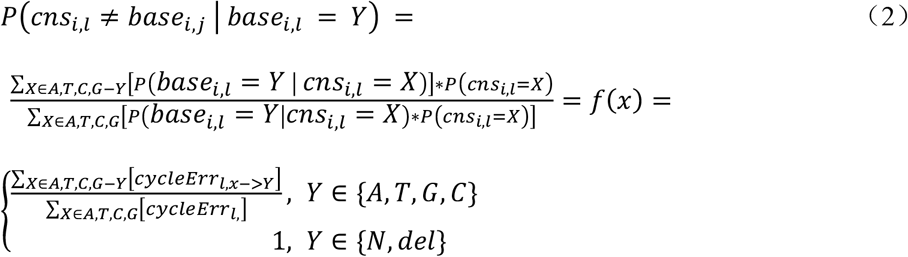

Based on the *l*-th base quality value of the *i*-th read (represented by *qual*_*i,l*_), the probability of sequencing errors *seqErr*_*i,l*_ on the current base can be calculated according to the formula (3):

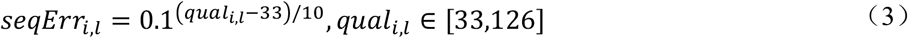

We use the error profile to correct the quality score of the base. The calculation method of the corrected sequencing error probability 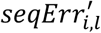 is as follows (4):

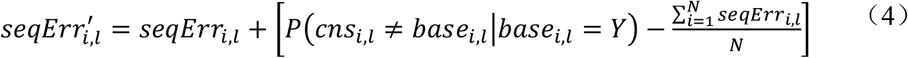

Finally, error probability 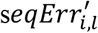 is converted to Phred quality score, and each read will get a new Phred quality score *qual*′_*i,l*_.

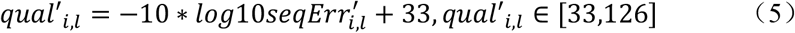

For a UMI cluster, the calculation of the consensus sequence requires the first alignment with the CIGAR information of each sequence, and then the posterior probability of different bases on each cycle of the consensus sequence is calculated. Finally, the base with the largest posterior probability is the cycle corresponding to the current consensus base.

The observation data at the *l*-th position of the *i*-th read is represented by *obs*_*i,l*_. It includes base, Phred quality score of the base, and genome alignment format:

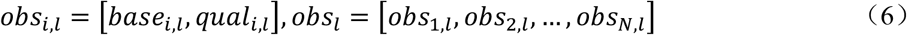

The posterior probability of the *l*-th base *cns*_*l*_ = *X* occurring in the consensus sequence was calculated based on *obs*_*i,l*_ according to Bayesian formula:

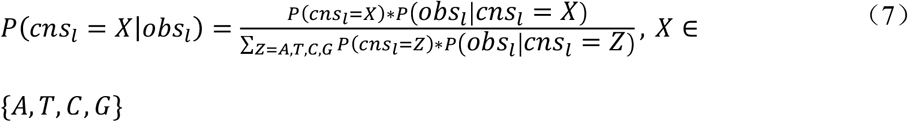

The prior distribution *P*(*cns*_*l*_ = *X*) can be calculated by the variation conservation score of the genome, and the prior probability is calculated from the conservation scores of the 45 vertebrate species compared with the human genome provided by the phyloP database. Denote the base at the *t*-th position of the reference genome as *ref*_*t*_ ∈ {*A, T, G, C*}, and the conservation score of germline variation at the *t*-th position is *mutS*_*t*_, then the probability that the *t*-th base is consistent with the reference genome is formula (8):

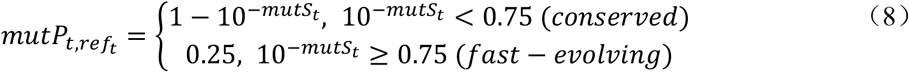

Since ∑_*M*∈{*A,T,G,C,del,ins*}_ *mutP*_*t,M*_ = 1, the probability of observing base *X* at position *t* in the genome after alignment can be calculated by the formula (9)

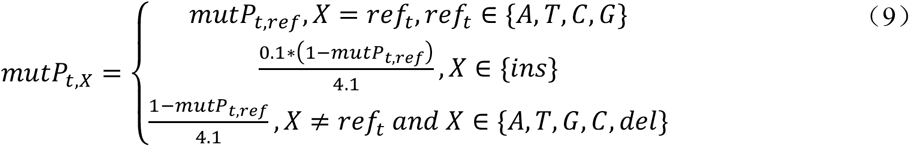

Based on the alignment position, the prior distribution of the bases in the *l*-th cycle of the sequences in this UMI cluster is *P*(*cns*_*l*_ = *X*) = *mutP*_*t,X*_. The conditional probability obeys the joint probability distribution based on the quality value of the sequence base. The calculation method is formula (10), where *Err*_*del*_ is the probability of deletion occurring:

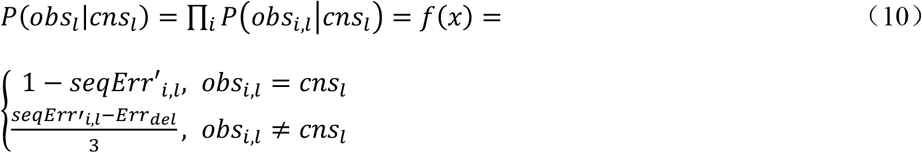

We set the probability threshold to 0.6. If the maximum posterior probability is greater than the threshold, the base with the maximum posterior probability is taken as the base corresponding to the consensus sequence. Otherwise, *N* is taken as the base corresponding to the consensus.

### Identification of methylation modification in bisulfite-free methylation data

In the process of duplex merging of two consensus sequences, ReadsCleaner can process not only paired-end NGS UMI data but also identify bisulfite-free methylation NGS data. According to the principle of bisulfite-free library construction of 5-methylcytosine (5mC is finally converted into T) (Liu, et al., 2019). ReadsCleaner will traverse each base in the process of merging two consensus sequences. Once it finds that two bases in the same position are inconsistent, and one is C and another is T, or one is G and another is A, the base of the corresponding locations at the reference genome are C or G, respectively. The program will correct the T base to C or the A base to G and record the corrected signal. In this way, all methylated sequences are corrected back to their original state before methylation, and with the help of the realignment module, the data can be restored to the original state while retaining the methylation signal.

We defined a tag in the SAM file format specifically for recording 5-methylcytosine (5mC) modification in sequences XM:Z: [0-9]+ (([Zz]|[Zz]+)[0-9]+(([Zz]\^[Zz]+)[0-9]+)*. The tag is to record the strand of the sequence, the original base type of base modifications, and the probability of each modification listed in the MM tag being correct. The XM tag consists of two parts: the numeric part represents the length of the sequence without methylation, and the Z or z string represents the positive chain and the negative chain methylation, respectively. For example, a 100bp sequence has the following XM tag: XM:Z:10Z0z68Z19, which implies that no methylation occurs in the first 10bp, methylation occurs in the 11bp and 81bp of the positive chain, and methylation occurs in the 12bp of the negative chain. Compared with third-generations of sequencing SAM file labels MM/ML, XM is mainly used in 5mC NGS, which can visually display the methylation position of the current sequence and distinguish between positive and negative strands has been methylated. The main differences between MM/ML and XM tags are shown in the Table 3.

**Table 3.**
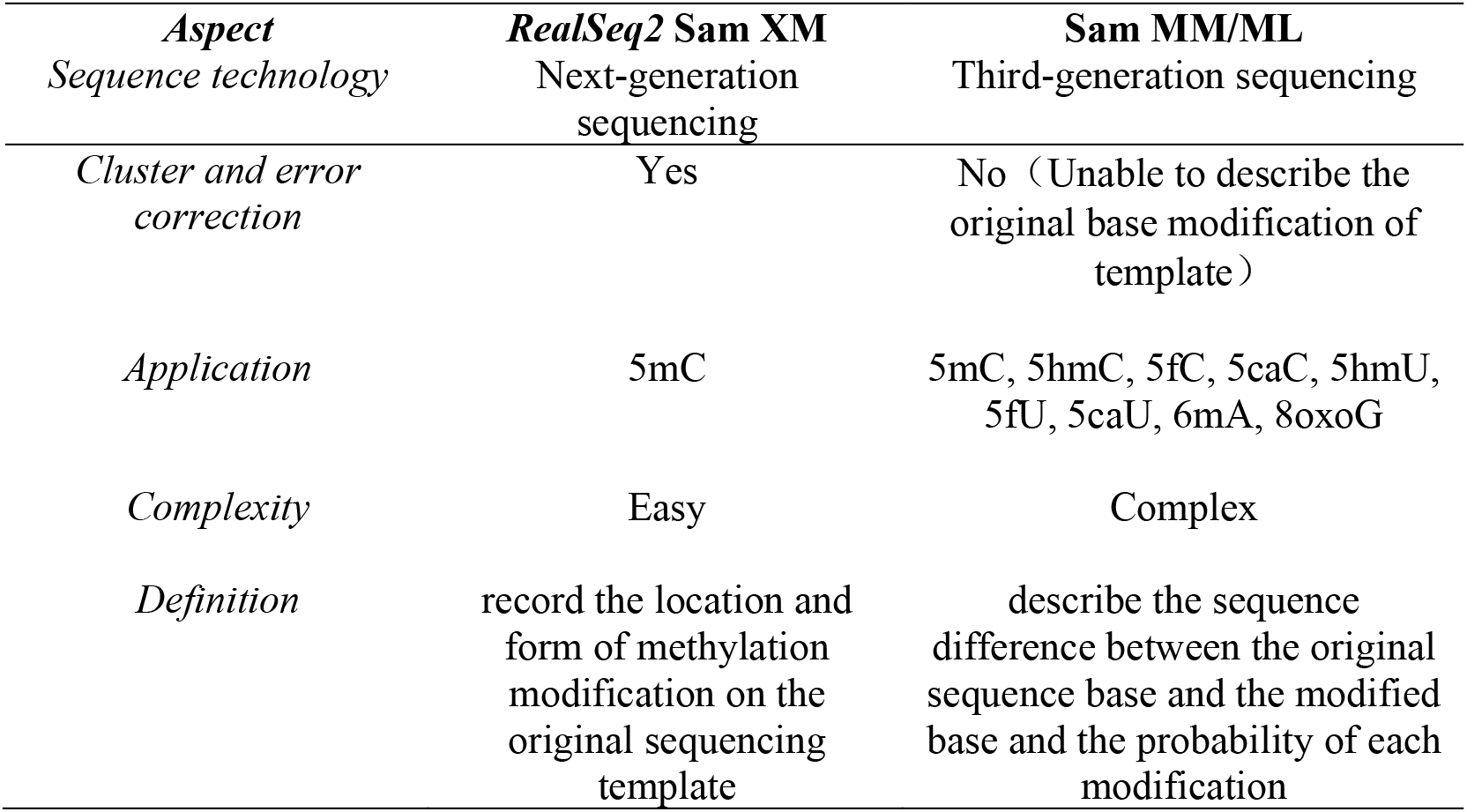
The difference between the *RealSeq2* XM tag and SAM MM/ML tag

| *Aspect* | *RealSeq2 Sam XM* | *Sam MM/ML* |
| --- | --- | --- |
| *Sequence technology* | Next-generation sequencing | Third-generation sequencing |
| *Cluster and error correction* | Yes | No (Unable to describe the original base modification of template) |
| *Application* | 5mC | 5mC, 5hmC, 5fC, 5caC, 5hmU, 5fU, 5caU, 6mA, 8oxoG |
| *Complexity* | Easy | Complex |
| *Definition* | record the location and form of methylation modification on the original sequencing template | describe the sequence difference between the original sequence base and the modified base and the probability of each modification |

Since the sequence of the highly methylated region will cause a large number of mismatches because of too many methylation positions. Some sequences will be aligned to more than one position and even cannot be mapped to the reference genome. ReadsRecycler submodule was designed to specifically handle these sequences. Instead of relying on sequence alignment information, ReadsRecycler can cluster sequences together through UMI tags. The sequence similarity is then used to further divide the sequence with a high degree of similarity into small UMI clusters. These small UMI clusters generate a consensus sequence based on the ReadsCleaner clustering error correction algorithm described above and then are proceeded with duplex merging. Finally, the realignment module filters out the sequence on which can be re-compared for output. With the ReadsRecycler module, data can be used to the maximum extent without waste.

## Results

We conducted the results of the RealSeq2 software in four aspects, including running speed, recognition of variable-length UMI error tolerance, error profile visualization, and downstream co-detection of CpG sites and mutations.

First, we performed UMI preprocessing on FASTQ files of three ∼5Gb samples, which were sequenced with 100bp paired-end by DNB-seq T7 sequencing platform, using RealSeq2, with or without UMI fault tolerance. The number of reads before UMI preprocessing, after UMI error-tolerant filtering, and after error-tolerant filtering was shown in Figure 4a. Clearly, as compared with a method of UMI processing which did not support error-tolerant, the method of using UMI error-tolerant processing can ensure the efficient use of data and reduce many reads that cannot be processed correctly due to UMI base errors.

**Figure 4.**
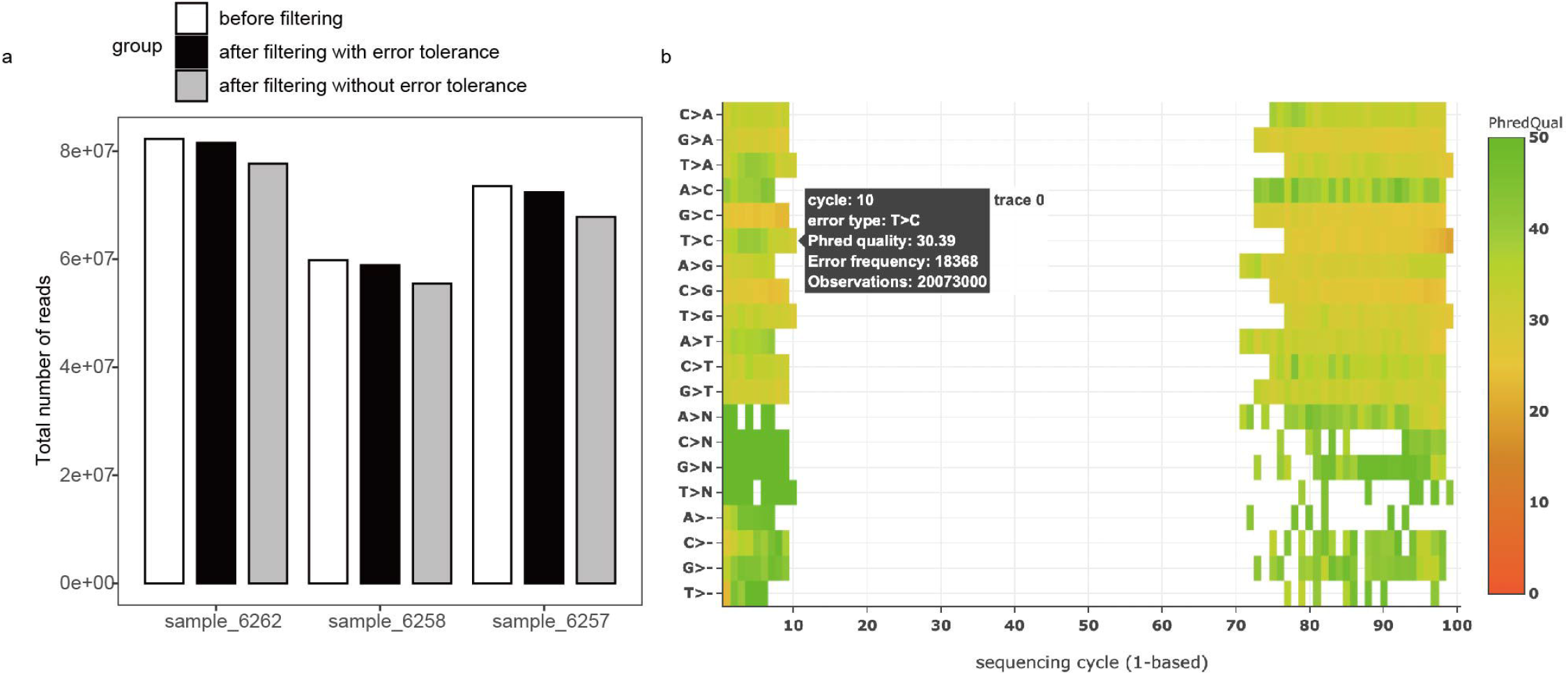
UMI preprocessing and error profiles; a) comparison of the total number of reads before filtering, after filtering with or without error tolerance; b) error profile of a test sample, total observations could be seen when you roll the mouse over the heatmap, among these, error type can indicate both real base and error base; Observations is the total number of bases observed at the cycle; error frequency is the number of error bases observed at the cycle; Phred quality = -10*log10 (Error frequency/Observations); The color from red to green indicates that the base quality value is from low to high.

*RealSeq2* can also visualize the base error profile of each sequencing cycle, display all quality control information of a sample in an HTML format, and dynamically view the detailed information of base errors of each cycle with mouse clicking. In Figure 4b, the 1 to 10 sequencing cycle on the left is the error rate of UMI statistics at the 5′ end, and the error rate tends to decrease but is not obvious, the 70 to 100 sequencing cycles on the right is the error rate of the UMI statistics at the 3′ end, and the error rate tends to increase. This is the same trend that the quality value of the subsequent cycle will deteriorate as the sequencing is measured.

Figure 5 is the IGV diagram after clustering and error correction (view as pair), a total of 174 reads are mapped on the 341,266 sites of chromosome 16, of which 107 reads support the base mutation C->T, 49 positive strands, 58 negative strands, reads without labeling bases at this position, may be either normal C or methylation C, but we can judge whether the site has been methylated through the XM tag. It is a powerful tool for supporting downstream methylation and mutation co-detection.

**Figure 5.**
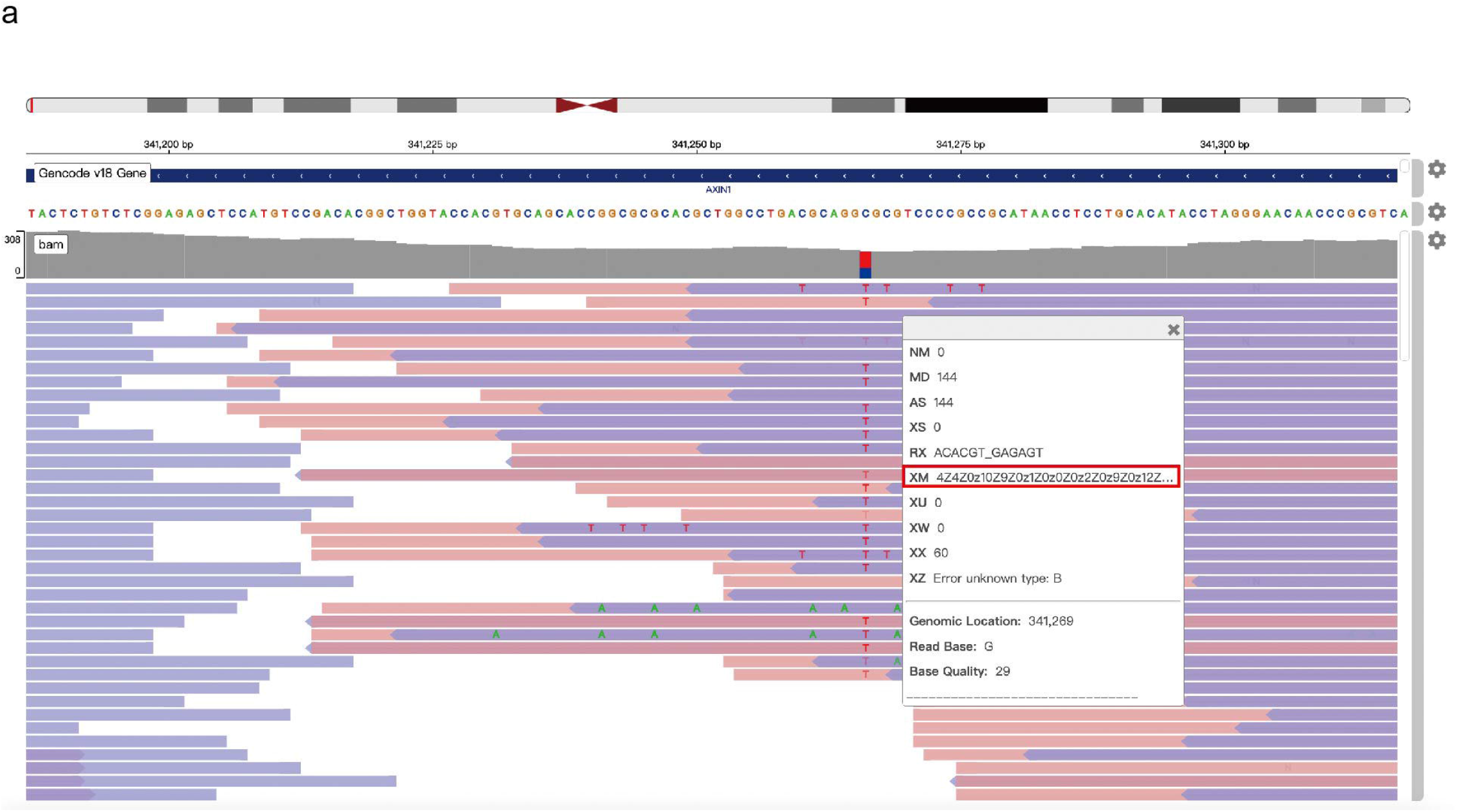
IGV diagram of BAM file with XM tag after clustering and error correction

## Discussion

In this work, we developed a comprehensive software called *RealSeq2* containing three modules: ReadsProfiler, ReadsCleaner, and ReadsRecycler. *RealSeq2* integrated sequencing quality control, UMI identification, sequence clustering and error correction based on UMI, which minimized the impact of PCR and sequencing errors on mutation detection. It also supported bisulfite-free methylation NGS data and defined an “XM” tag to record the methylation signal.

Compared to classic UMI trimming software such as *fastp*, the ReadsProfiler submodule not only supports read quality control and retains the advantage of fast preprocessing but also supports the identification of variable-length UMI and the UMI with base mismatch and deletion. Besides, ReadsProfiler can also estimate sequencing cycle error rate, generate sample quality control visualization reports, and support equal-length sequence output for better compatibility with downstream software. ReadsProfiler is equivalent to fastp in time consumption. We also reconstructed the sequence clustering and error correction algorithm using Bayesian Maximum A Posteriori based on Gencore basic structure and implemented multithreading. ReadsCleaner submodule greatly improves error correction accuracy and running speed compared to prevailing software such as Gencore and Fgbio. Importantly, it supports bisulfite-free methylation data apart from supporting the whole genomic sequencing data. As a new burgeoning methylation technology, bisulfite-free has obvious advantages over traditional bisulfite sequencing (Liu, et al., 2019). ReadsCleaner is such a submodule that can highly restore original CpG methylation site information in bisulfite-free methylation sequencing. We also defined a new tag XM in SAM file format to record methylation positions of the current sequence and distinguish the positive and negative strands. ReadsRecycler was specifically developed for bisulfite-free data to correct and realign a large number of sequences that cannot be aligned to the reference genome due to methylation to improve data utilization. In detail, The bisulfite-free data preprocessing by *RealSeq2* can retain both mutations and methylations, which is conducive to the simultaneous detection of mutations, CNVs, fusions, methylation, fragment allosteric. Especially for quantification of tissues origin (Loyfer, et al., 2023) and donor-derived cfDNA (dd-cfDNA) by methylation. In detail, donor-specific single-nucleotide polymorphisms (SNPs) was used to quantify dd-cfDNA in recipient plasma, which was a biomarker for organ injury (Jaikaransingh and Kadambi, 2021; Paul, et al., 2021). In conclusion, it is necessary to have an all-in-one software with high quality, fast speeding, and supporting bisulfite-free methylation data.

The limitation of *RealSeq2* is that it only supports paired-end UMI inputs and 1bp mismatch and deletion in UMI. In the follow-up work, *RealSeq2* will support single-end UMI reads, add the parameter of the maximum number of mismatches or deletions bases, and improve the running speed. Another limitation of this study is that the accuracy of methylation sites and mutation sites was tested using simulated data. We will test and improve the accuracy of the software with the data of known methylation sites in the following study. The *RealSeq2* software currently only supports the analysis of bisulfite-free libraries, but the principle of distinguishing mutations and methylation is also applicable to BS-seq libraries.

In conclusion, *RealSeq2* is the preferred upstream analysis software for the co-detection of ultra-low frequency mutations and bisulfite-free methylation data. The error profile provides data support for downstream analysis. Additionally, XM tags will become a standard protocol for recording methylation signals.

## Key Points

- We developed RealSeq2, a pipeline for the data pre-processing of cfDNA sequencing raw data in liquid detection. It contains functions of adapter removal, quality control, UMI identification, generating consensus reads by clustering and error correction, storing information of methylation modifications by defining a new BAM tag XM, and providing support for the downstream analysis of methylations and mutations co-detection.
- Compared to fastp software and gencore, RealSeq2 supports the recognition of variable-length UMI, the visualization of error profiles of each sequencing cycle, and improves running speed.
- The clustering and error correction process of RealSeq2 adopts the principle of Bayesian posterior probability. The whole process uses the error profile to correct the base quality value of a sequence which represents the probability of an error and uses the method of maximum posterior probability to calculate the base form of each cycle on the sequence by cycling through all sequencing cycles. Finally, the posterior probability of the base is used as the base quality value.
- RealSeq2 supports the ability to detect methylations and mutations simultaneously in one bisulfite-free methylation sequencing.

## List of abbreviations

UMI: unique molecular identifier
5mC: 5-methylcytosine
WGBS: Whole Genome Bisulfite Sequencing
VAF: variant allele frequency

## Declarations

### Ethics approval and consent to participate

Not applicable

### Consent for publication

All study participants provided informed consent, and the study design was approved by an ethics review board.

### Availability of data and materials

All data generated or analyzed during this study are included in this published article, the executable program of RealSeq2 is https://hub.docker.com/r/geneplus/realseq2

### Authors’ contributions

TL, YG, XX, and XY envisioned this project. TL and HF developed RealSeq2 algorithm, implemented the project, and conducted the analyses; MS, ZL, and XG wrote the manuscript. ML, TC, KW, and EY helped in analyzing the data. All authors have read and approved the final manuscript.

## Acknowledgements

Thanks to all authors for their contributions to the software and the article.

